# ATP and glutamate coordinate contractions in the freshwater sponge *Ephydatia muelleri*

**DOI:** 10.1101/2024.05.23.595635

**Authors:** Vanessa R Ho, Greg G Goss, Sally P Leys

## Abstract

Sponges (phylum *Porifera*) are an early diverging animal lineage that lacks both conventional nervous and muscular systems, and yet they are able to produce coordinated whole-body contractions in response to disturbances. Little is known about the underlying signaling mechanisms in coordinating such responses. Previous studies demonstrated that sponges respond specifically to neuroactive chemicals such as L-glutamate and γ-amino-butyric acid (GABA), which trigger and prevent contractions respectively. Genes for purinergic P2X-like receptors are present in several sponge genomes, leading us to ask whether ATP works with glutamate to coordinate contractions in sponges as it does in other animal nervous systems. Using pharmacological approaches on the freshwater sponge *Ephydatia muelleri*, we show that ATP is involved in coordinating contractions. Bath applications of ATP cause a rapid, sustained expansion of the excurrent canals in a dose-dependent manner. Complete contractions occur when ATP is added in the presence of apyrase, an enzyme that hydrolyzes ATP. Applying ADP, the first metabolic product of ATP hydrolysis, triggers complete contractions, whereas AMP, the subsequent metabolite, does not trigger a response. Blocking ATP from binding and activating P2X receptors with pyridoxalphosphate-6-azophenyl-2’,4’-disulfonic acid (PPADS) prevents both glutamate- and ATP-triggered contractions, suggesting that ATP works downstream of glutamate. Bioinformatic analysis revealed two P2X receptor sequences, one which groups with other vertebrate P2X receptors. Altogether, our results confirm that purinergic signaling by ATP is involved in coordinating contractions in the freshwater sponge suggesting a role of ATP-mediated signaling that predates the evolution of the nervous system and multicellularity in animals.

**Summary statement:** Nerveless sponges coordinate a sneeze-like reflex using glutamate and ATP signaling to expel water from the body.

## Introduction

The ability to sense and respond to environmental cues is essential for all forms of life. Most animals use the nervous system for this, but sponges (Porifera) and many multicellular non-animal organisms, e.g., the Venus flytrap (*Dionaea muscipula*) and the aggregating slime mold *Dictyostelium discoideum* (Amoebozoa), carry out sensation and coordination in without it. The means by which signals are transmitted between cells to coordinate these activities is as diverse as the organisms themselves, with some quite well known, such as the chloride potential of *Dionaea* (Hedrich and Kreuzer, 2023; Hodick and Sievers, 1988) and cAMP signaling in *Dictyostelium* (Miura and Siegert, 2000; Swanson and Lansing Taylor, 1982), but in others, such as sponges, signaling mechanisms in coordinating behaviour are still unknown.

Sponges are an ancient metazoan lineage (Schultz et al., 2023). Typical animal features such as bilateral symmetry and conventionally defined organ systems (e.g., nervous and muscular systems) are absent in sponges. Despite the lack of a nervous system, it is known that all sponges sense and respond to external stimuli (de Ceccatty, 1974; Elliott and Leys, 2007; Nickel, 2004; Parker, 1910). As sessile filter feeders, sponges are susceptible to clogging their aquiferous canal system with unwanted debris that enters alongside food particles. To address this problem, sponges can control their feeding current – how this occurs depends on the physiological characteristics of the sponge itself. The syncytial tissues of glass sponges (Hexactinellida) (Mackie et al., 1983) propagate electrical currents to coordinate the rapid arrest of the feeding current in response to disturbances (Leys, 2003; Leys and Mackie, 1997; Leys et al., 1999; Tompkins-MacDonald and Leys, 2008). On the other hand, multicellular sponges (Demospongiae, Calcarea, and Homoscleromorpha) respond by shutting the incurrent pores (ostia) and constricting the aquiferous canal system to halt the feeding current (Leys and Meech, 2006).

Whole-body contractions are a type of behaviour displayed by sponges that function like a sneeze to expel wastes and other irritants from the internal canal system (Elliott and Leys, 2007; Kornder et al., 2022). How contractions are coordinated is not well understood: they are too slow to be propagated by electrical impulses, and gap junctions have not yet been identified (Leys, 2015). It was once thought that signal propagation occurred by physically connected cells pulling on each other (Parker, 1910; Pavans de Ceccatty et al., 1960), but this is unlikely because sponges continue contracting despite having torn their tissues (Ludeman et al., 2014). At present, physiological and molecular evidence suggest coordination is accomplished by a paracrine signaling system that may have been a precursor to the nervous system (Fields et al., 2020; Kenny et al., 2020; Musser et al., 2021).

Sponges respond to neuroactive substances such as L-glutamate and γ-amino-butyric acid (GABA) (Elliott and Leys, 2010; Ellwanger et al., 2007), among others (reviewed in Leys, 2015). Glutamate and GABA are the most abundant excitatory and inhibitory neurotransmitters in vertebrate nervous systems respectively (Andersen and Schousboe, 2023). The former triggers contractions while the latter prevents them in the freshwater sponge *Ephydatia muelleri* (Elliott and Leys, 2010).

In many animal nervous systems, glutamate signaling often works in concert with ATP signaling (Gu and MacDermott, 1997; Hamilton et al., 2008; Rimbert et al., 2023; Shen et al., 2017). As the main energy currency of cells, ATP is both abundant (2-8 mM in mammalian cells; Traut, 1994) and highly regulated (Dunn and Grider, 2023). These characteristics make it a readily available multi-purpose molecule for fueling cellular activities and acting as an intercellular chemical messenger (Fountain, 2013; Novak, 2003; Rimbert et al., 2023). Purinergic (ATP) signaling is essential for the function of various tissues, and is the most prominent form of communication between glia and neurons (Di Virgilio et al., 2023). ATP in excess of normal cytosolic concentrations is stored in secretory vesicles with other chemical messengers (Bonora et al., 2012; Novak, 2003). In the nervous system, it is co-released with other neurotransmitters (Fellin et al., 2006) in response to membrane stretching, glutamate transmission, or even by ATP signaling itself (Fields and Burnstock, 2006). Activation of the P2X receptor (P2XR) channels by extracellular ATP allows cations, namely Na^+^, K^+^, and Ca^2+^, to pass into the cell (North, 2016). This in turn leads to a complex cascade of downstream effects including, but not limited to, Ca^2+^ signaling (Hamilton et al., 2008), glutamate release (Nakatsuka et al., 2003) and subsequent auto- and paracrine functions (Hinoi et al., 2004), and more ATP release (Shen et al., 2017).

The nervous system-like characteristics of coordination in multicellular sponges led us to ask whether purinergic signaling by adenosine triphosphate (ATP) plays a role in coordinating the behaviour of sponges. We took advantage of the ease of laboratory culture and transparent tissues of the freshwater species *Ephydatia muelleri* (Demospongiae; Lieberkühn, 1856). Using a pharmacological approach, we test the hypothesis that sponges respond to exogenous ATP in a concentration-dependent manner, and that ATP signaling occurs downstream of glutamate signaling. We also carried out a phylogenetic study of sponge P2X receptors, examining the evolutionary relationships of P2XRs in metazoans and non-metazoans. The evidence we provide from both pharmacological and bioinformatic approaches bridges a gap in knowledge in the physiology of sponges, addressing the question of how cells communicate across the body of the sponge to coordinate behaviour.

## Materials and methods

### Sponge collection and culture

Fragments of *E. muelleri* containing gemmules were collected in the winter months of 2020 and 2021 from O’Connor Lake, British Columbia, Canada. These were stored in unfiltered lake water in the dark at 4°C in plastic bags, aerated monthly, until use. Cleaning and culturing were done as described previously (Leys et al., 2019). Briefly, gemmules were mechanically dissociated from the dead sponge skeleton in ice cold water and sterilized with 1% hydrogen peroxide. Sponges were cultured on flame-sterilized 22 mm^2^ glass coverslips in ~20 mL Strekal’s medium (Strekal and McDiffett, 1974) in 60 mm diameter Petri dishes and kept in the dark. Fifty percent water changes were done every 2-3 days until 6-10 days post-hatching (Stage 5). Sponges with a fully formed and functional aquiferous canal system and a single, upright osculum (Fig. 1) were used for experiments.

**Figure 1.**
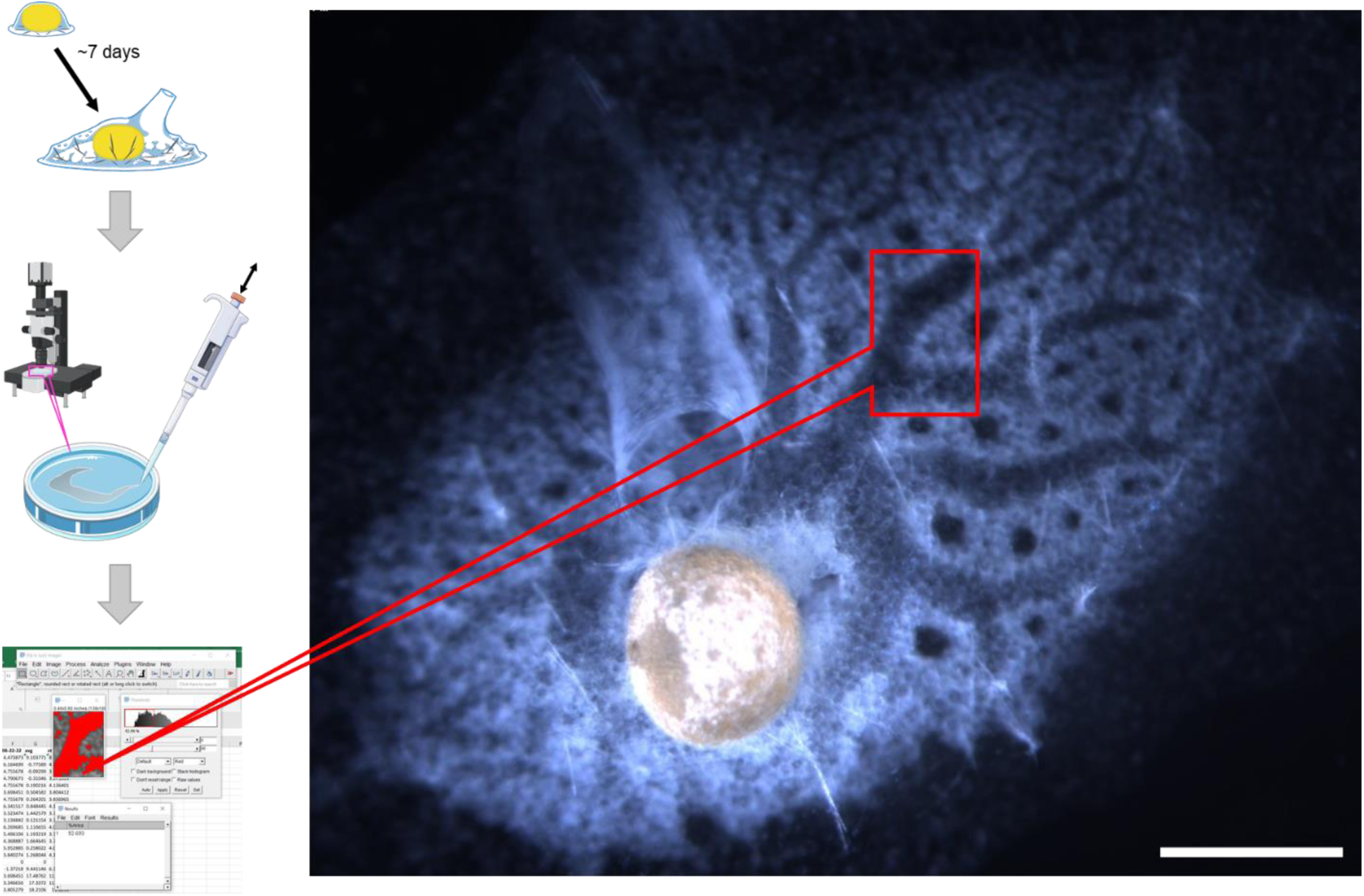
Overview of the experimental workflow and top view of a representative stage 5 juvenile sponge, *Ephydatia muelleri*. After hatching and culturing sponges, time-lapse images of contraction responses to mechanical (not shown) or chemical stimulation were taken on a stereomicroscope. Image analysis was used to measure a selected segment of the excurrent canal system and processed to determine the average relative change in canal area. Scale bar, 0.5 mm.

### Test substance preparation and application

Stock solutions of pharmacological substances were prepared with Milli-Q water (Millipore-Sigma) in the following concentrations: 100 mM adenosine 5’-triphosphate (ATP) disodium salt hydrate (Sigma-Aldrich A2383), 20 mM L-Glutamate (Sigma-Aldrich G1251), 100 mM PPADS (Sigma-Aldrich P178), and 200 units/mg apyrase (Sigma-Aldrich A6410). ADP and AMP were obtained from G. Goss.

Prior to filming, the culture medium was removed until 10 mL was left in the Petri dish. The sponges were then carefully set on the microscope stage to prevent triggering a premature contraction. All sponges were filmed for 5 min prior to the application of any substance to ensure no contraction was occurring. If sponges were already in the middle of a contraction, they were left alone for at least 1 h to allow the completion of the contraction before proceeding with the experiment.

Dilution of test chemicals was calculated to a final volume of 10 mL in the bath application, and was mixed with the previously withdrawn medium from the Petri dish to make 1 mL of 10X solution. This was then gently mixed around the dish by gently pipetting 7-8 times, ensuring sponges were fully and evenly exposed to the test substance. To validate the mixing technique, fluorescein dye (134 mg/L 0.2 µm filtered twice) was used as a visual equivalent (not shown). For a negative control, 1 mL of the withdrawn Strekal’s medium was mixed back in to the Petri dish. Sponges treated with multiple substances (apyrase and PPADS experiments) were first bathed in the indicated drug and left undisturbed for 20 min prior to the application of glutamate or ATP.

### Digital time-lapse and data acquisition

Sponges were viewed with a stereomicroscope (Olympus SZX-12) and digital time lapse images were captured with a QI-Cam colour CCD camera (Retiga QImaging) and Northern Eclipse version 7 (Empix Imaging, Toronto). Images were saved as 8-bit grayscale except for PPADS experiments, which were captured in 16-bit colour for splitting colour channels. Single-treatment time lapse images were captured at 20 s intervals for 60 min, dual-treatment time lapses were captured at 30 s intervals for 90 min. Behavioural responses were analyzed and quantified using FIJI ImageJ version 1.53t (Schindelin et al., 2012) by measuring changes in the excurrent canal area during a contraction (Ho and Leys, 2024). Canal sections undergoing the most visually distinguishable changes in size during the course of a contraction were selected and measurements were taken as percent area fractions. The data was exported to MS Excel (2016) to calculate the mean relative change in area after application of stimulus. Graphs were generated in MS Excel and figures were processed in Adobe Illustrator CS5.

### Bioinformatic analysis and phylogenetic tree building

The protein sequences for P2X-like receptors in *E. muelleri* were obtained by identifying the most highly expressed genes from RNAseq differential transcript expression data (Kenny et al., 2020); available on EphyBase (https://spaces.facsci.ualberta.ca/ephybase/). The differential transcript expression values of P2XRs in *E. muelleri* showed low relative transcript abundance. We selected the two most highly expressed genes of the group: Em0004g77a and Em0004g666a (FPKM 7.37 and 3.15 respectively, Dataset 1). The corresponding protein sequences were obtained from functional annotation results (Kenny et al., 2020) also on EphyBase, and verified by performing a BlastP search (National Center for Biotechnology Information; NCBI).

P2X receptor sequences from diverse metazoan and non-metazoan lineages were collected by searching the NCBI protein database (Dataset 2). Additional sponge P2X-like receptors (*Spongilla lacustris, Aphrocallistes vastus, Sycon coactum,* and *Eunapius fragilis*) were identified by tBLASTn against their respective transcriptomes (Riesgo et al., 2014; Windsor Reid et al., 2018); data available on https://leyslab.weebly.com/data-available.html). Protein sequences <200 amino acids were excluded.

Sequences were aligned with MAFFT version 7 (Katoh et al., 2017) and trimmed with TrimAl v1.3 (Sánchez et al., 2011). SeaView version 5 (Gouy et al., 2021) was used to visualize the aligned sequences and fragmented or poorly aligned sequences were removed manually. The phylogenetic tree was computed on IQ-Tree using 1000 ultrafast bootstrap approximations (Hoang et al., 2017), and the resulting consensus tree was viewed with Fig Tree v1.4.4 (Rambaut, 2018) and rooted with *Dictyostelium discoideum* P2X sequence.

The predicted 3D structures of selected *E. muelleri* P2XR subunits, Em0004g77a and Em0004g666a, were compared to the well-characterized structure of the *Danio rerio* P2X4 subunit (NCBI NP_945338.2), all of which were generated on Phyre2 (Kelley et al., 2015) and visualized on EzMol (Reynolds et al., 2018). Consensus sequence shading of phylogenetic tree P2XR sequences was made with Boxshade (Hofmann et al., 1997) at 0.75 sequence agreement (range 0-1) (Fig. S1).

## Results

### Variations in contraction responses to mechanical or chemical stimulation

Sneezes triggered by endogenous signals (i.e. without application of chemicals) were induced by shaking Petri dishes vigorously, which provided a reference for how excurrent canals change in size, first expanding and subsequently contracting (Movie 1). We follow previous researchers in referring to this as a ‘sneeze’ (Colgren and Nichols, 2022; Elliott and Leys, 2007; Ludeman et al., 2014) and in our work we record the dilation of the excurrent canals (ECs) as the expansion phase of the sneeze, and the constriction of the ECs as the contraction phase of the sneeze. It is generally considered that the opposite reaction is occurring in the incurrent canals (ICs) during this process. To trigger a sneeze pharmacologically, we added 70 µM L-glutamate (Movie 2). In shaken sponges, expansion of excurrent canals start at one end of the sponge near the osculum, and move in a wave-like manner across the body of the sponge, whereas pharmacological stimulation caused all canals to respond simultaneously (movies). In both cases, the percent area fraction showed excurrent canals expanding and constricting, together the events making one “contraction cycle” (Fig. 2). Mixing the medium with pipetting did not trigger sponge contractions (Movie 3).

**Figure 2.**
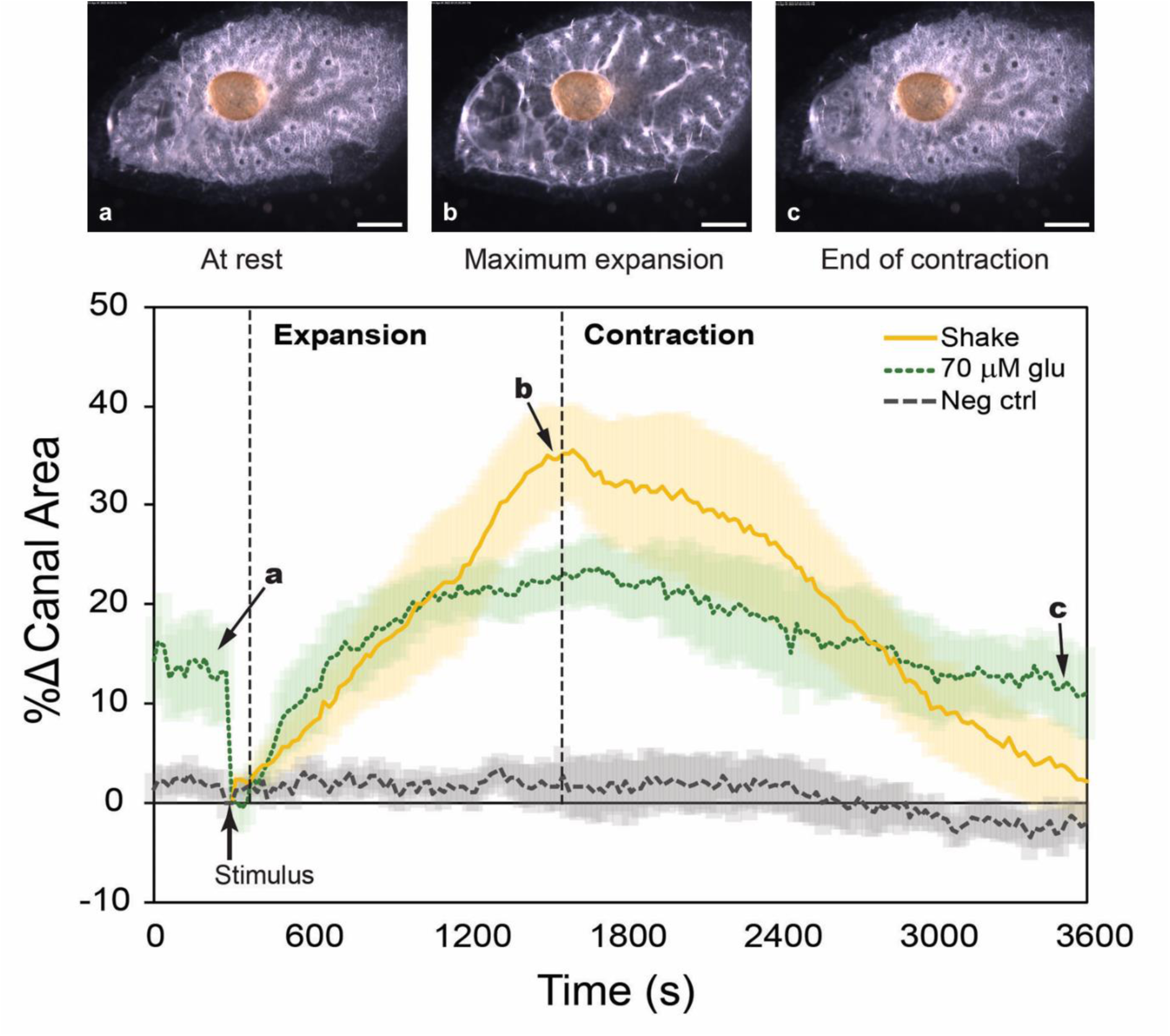
Progression of one contraction cycle in response to mechanical or chemical stimulation. Images show the stages a contraction in sponges over the course of a contraction (scale bar, 0.5mm). Labels (a,b,c) and arrows on the graph indicate the corresponding time and canal size of stage in the contraction shown by the images. Changes in excurrent canals in response to stimulation are plotted as follows: physical shaking (solid gold line), 70 µM L-glutamate (dotted green line), or negative control (dashed grey line). n=7 individuals for each treatment, shading behind plotted lines indicate the standard error of the mean.

### ATP treatment

Sponges responded to ATP in a dose-dependent manner (Movies 4-7). Application of lower doses of ATP (20 or 40 µM) caused a small constriction of the excurrent canals (EC) (Fig. 3A). At the higher concentration of 40 µM ATP, ECs eventually expanded slightly (<10%) before relaxing to the initial resting size, whereas 20 µM ATP caused a slight constriction of EC and slow relaxation, but no expansion. High concentrations of exogenous ATP (100 and 200 µM) caused the rapid and intense dilation of ECs and prevented their constriction. These dosages triggered responses so strong that some sponges tore the ‘tent’ which forms the outer tissue of the sponge. At 200 µM ATP, expansions of the ECs were faster and more intense than with 100 µM ATP (Fig. 3A) with more sponges tearing their surface tissues than those exposed to lower concentrations. The most severe responses caused the aquiferous canals themselves to tear (Movie 4).

**Figure 3.**
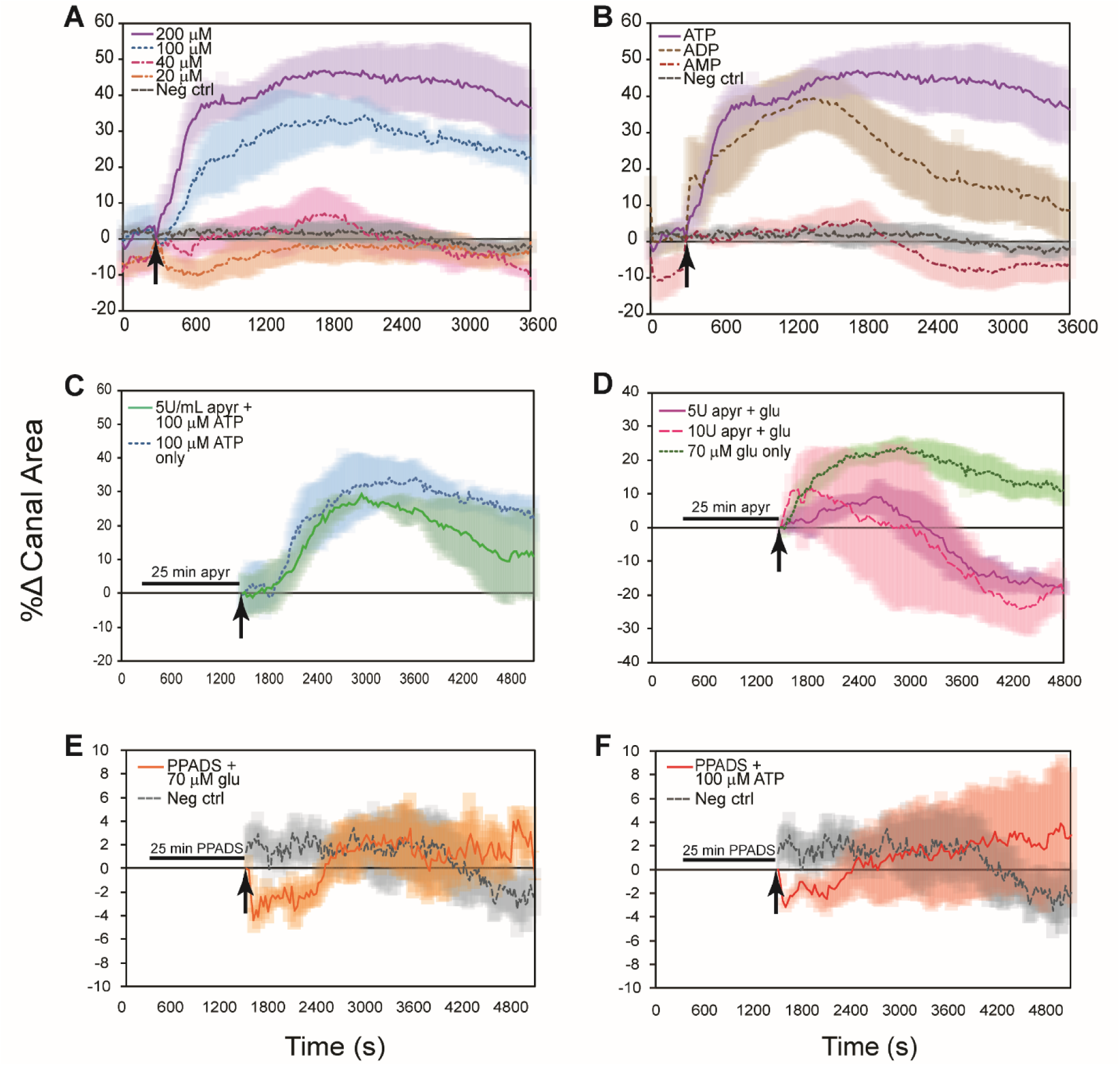
Pharmacological treatments have different effects on the contraction response. (A) Responses to ATP are biphasic and concentration-dependent. While 20 and 40 µM ATP cause a temporary constriction of excurrent canals, 100 and 200 µM ATP trigger the rapid expansion of ECs but prevents the constriction phase to complete a contraction cycle. (B) Sponges respond to 100 µM of ATP and ADP, but not AMP. (C) Addition of 5U/mL apyrase to the dish medium allows for a complete contraction when triggered by 100 µM ATP. (D) Addition of apyrase causes the excurrent canals to become constricted when 70 µM L-glutamate is added. (E) & (F) PPADS blocks either glutamate- or ATP-induced contractions from occurring. n=7 individuals for each treatment in panel A, n=5 in panels B-F (n=4 for 10 U/mL apyrase + glutamate, panel D). Shading behind plotted lines represents the standard error of the mean.

### Pre-treatment with the hydrolytic enzyme apyrase

Whereas treatment with 100 µM ATP caused the EC to remain expanded, pretreatment with 5U/mL apyrase allowed 100 µM ATP-treated sponges to complete a full expansion and contraction cycle (Fig. 3C, Movie 8). Treatment with 70 µM glutamate after pre-incubation in apyrase caused sponges to rapidly expand and contract their ECs, and by the end of the experiment, the ECs were more constricted than at the beginning of the experiment (Fig. 3D, Movie 9). Increasing the apyrase concentration to 10U/mL caused a greater variability in the responses, and some sponges sneezed in response to the apyrase itself. At the end of the experiment, the ECs of sponges treated with 10U/mL apyrase were more constricted than those of sponges pre-treated with 5U/mL apyrase (Fig. 3D, Movie 10). Sneeze responses (expansion and constriction of the ECs) in sponges pre-treated with apyrase were more intense than those in sponges treated with glutamate only, and the contractions travelled in a wave-like manner from one end of the sponge body to the other. However, when ATP was added in the presence of apyrase, the tissues did not tear apart as they did when only treated with ATP.

Furthermore, apyrase itself appeared to have an effect on the ECs. In all experiments, sponges were observed to respond when the apyrase was added into the Petri dish (Movies 8-10). Sponges briefly expanded and contracted their ECs, while some did the opposite by constricting then expanding the canals. This occasionally triggered a sneeze that propagated across the body of the sponge in a wave-like manner similar to sneezes triggered by mechanical stimulation.

### Treatments with ADP and AMP

Sponges responded rapidly to treatment with ADP, and the amplitude of the expansion phase (EC dilation) was nearly identical to responses triggered by the equivalent concentration of ATP (Fig. 3B, Movie 11). However, a significant difference was that ADP triggered a complete contraction cycle, while the same concentration of ATP immobilized the ECs in an expanded (dilated) state. Contractions triggered by either ATP or ADP were vigorous, but none of the sponges stimulated by ADP tore their tissues. Sponges did not respond at all to AMP (Fig. 3B, Movie 12).

### Pre-treatment with PPADS

Sponges treated with 100 µM PPADS could not be stimulated to sneeze with 70 µM glutamate (Fig. 3E, Movie 13) or 100 µM ATP (Fig. 3F, Movie 14). The activity of the ECs was no different from that of the negative controls, with only subtle and gentle movements.

### Bioinformatic analysis of purinergic receptors

P2XRs were found in all branches of Unikonta in the eukaryotic domain of life (Fig. 4A). Rooted to *D. discoideum* P2X, our tree showed two distinct branches of P2X receptors: one with only invertebrates, and another that included both vertebrate and invertebrate P2XRs (Fig. 4B). The group containing both vertebrate and invertebrate P2X-like receptors shared strong sequence consensus both in the characteristic motifs of P2X receptors (North, 2002), and in sharing many amino acid sequences along the length of the protein (Fig. 4B; Sup. 3).

**Figure 4.**
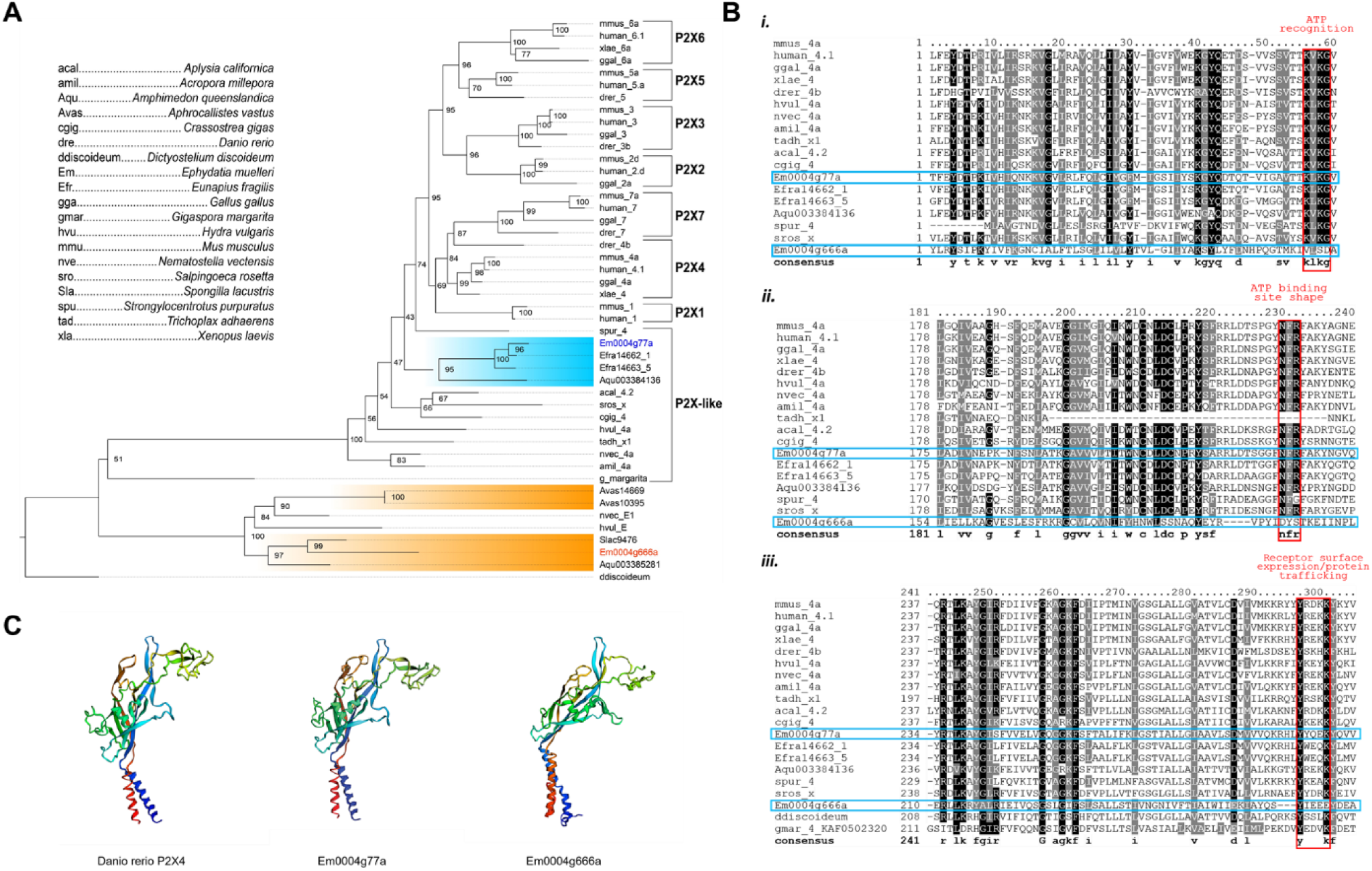
Bioinformatic analysis of *E. muelleri* P2X receptor sequences. (A) Consensus tree of evolutionary relationships of metazoan and non-metazoan P2X and P2X-like receptor amino acid sequences, rooted to the aggregating slime mold *Dictyostelium discoideum*. Phylogeny was generated in IQ-TREE v1.6.12, branch support values were calculated using 1000 ultrafast bootstrap replicates shown as number values (0-100) in nodes. Sponge sequences are highlighted and *E. muelleri* sequences are labeled in blue (Em0004g77a) and red (Em0004g666a). Branch lengths are not representative of time scales. (B) Boxshade multiple sequence alignment of one P2X receptor sequence from each organism, sequence agreement fraction set to 0.75 and consensus sequence text is bolded. Only sequences with 100% bootstrap support values in relation to vertebrate P2X sequences are shown (except in panel *iii*). The complete set of shaded sequences is available in Supplemental data 3. P2X conserved motifs are outlined in red and are listed from N-C along the protein sequence: i) KxKG ii) NFR iii) YxxxK. (C) 3-dimensional models of P2X receptor subunits of *Danio rerio* (RefSeq NP_945338.2) and *E. muelleri* show highly similar predicted protein structures, though Em0004g666a was not found to have the characteristic P2X conserved motifs in its amino acid sequence.

Predicted 3D constructions of the receptor subunit showed structural similarities between both *E. muelleri* sequences and a well-characterized P2XR from a zebrafish, all sharing the so-called ‘dolphin’ shaped structure that is characteristic of vertebrate P2X receptors (Fig. 4C). Em0004g77a shared conserved amino acid sequences with well-studied vertebrate P2XRs. In contrast, Em0004g666a lacked the consensus sequences for ATP binding and recognition sites, and only had a partial match to the convserved protein trafficking sequence KxKG (Fig. 4Biii). However, the trafficking sequence was present in amoeba (*D. discoideum*) and fungus (*Gigaspora margarita*) P2X sequences (Fig. 4Biii).

## Discussion

The signaling pathways that coordinate behaviour in sponges parallel many of the specialized physiological systems of vertebrates and are highly conserved in the evolutionary history of animals. Our data show that ATP signaling is involved in coordinating the contraction behaviour in sponges because sponges respond to ATP in a concentration-dependent manner. Moreover, our data indicate that purinergic signaling by ATP is necessary for triggering the expansion phase of the sneeze response and that ATP works together with glutamate signaling to coordinate the contraction phase of the sneeze behaviour. The balance of these chemical signals is important for propagating the contraction across the tissues to expel fluid from the sponge aquiferous system.

### ATP signaling is involved in coordinating contractions

We first investigated whether ATP would induce a response in sponges, and how it might affect the sponge sneeze behaviour. The transcriptome of *E. muelleri* contains several P2XRs (Kenny et al., 2020) (Supplemental data 1). As with most cellular receptors, P2XR subtypes each have their own ligand-binding affinities (North, 2002). Our data show that sponges responded in a concentration-dependent manner to ATP, suggesting different subsets of P2XRs were activated and caused different effects.

Part of our initial hypothesis was that ATP acts downstream of glutamate. Since glutamate can trigger a sneeze (Elliott and Leys, 2010), we thought glutamate caused the expansion of ECs in the first half of the expansion-contraction cycle, and ATP was responsible for ending the contraction by constricting ECs. We therefore expected to see the ECs constrict in the presence of exogenous ATP, essentially quenching the sneeze behaviour. However, while sponges do have a specific ATP-induced response, we found that with enough ATP, the ECs expanded and were immobilized in this dilated state instead.

Our data suggest ATP induces a biphasic response: at high doses of ATP (100 and 200 µM), ECs expand vigorously, and at lower concentrations (20 and 40 µM) they appear to constrict. At 40 µM ATP, the ECs first constrict before gradually expanding and contracting, the latter resembling a slow, smaller sneeze-type behaviour (Fig. 3A). This suggests that there is a threshold for ATP-induced activation of the activities that trigger the start of a sneeze.

### ATP and glutamate have different roles in coordinating contractions

In the vertebrate nervous system, there are two ligand-gated receptor families that mediate postsynaptic cellular activities: ionotropic receptors and G-protein-coupled (metabotropic) receptors (Grubbs, 2008). Whereas P2X receptors are ionotropic, allowing direct influx of ions into the cell when activated by ATP, metabotropic glutamate receptors involve several downstream steps to transduce chemical signals (Hamilton et al., 2008). Our experiments showed that hydrolysis of extracellular ATP by apyrase modulated the pharmacologically stimulated sneeze behaviour. While adding ATP alone caused the excurrent canals to expand and remain expanded, when ATP was added to the sponge in the presence of apyrase, the sponge was able to complete the full expansion-contraction cycle (the sneeze).

In vertebrates, after ATP is released by cells, the purinergic signal is terminated by degradation of ATP by ectonucleotidases (Novak, 2003). When adding a high concentration of ATP to sponges, a sneeze is triggered, but it appears as though the receptors are overstimulated and the ATP overwhelms endogenous removal mechanisms, thus causing the sustained expansion of ECs. Therefore, pre-incubation with the ectonucleotidase apyrase aids in the breakdown of ATP and ends the otherwise sustained signal transmission from excess ATP (Del Campo et al., 1977).

Individual sponges reacted differently to apyrase, resulting in varied responses such as premature contractions (data not shown). Apyrase is highly efficient at metabolizing ATP (Del Campo et al., 1977), and may be metabolizing endogenous ATP the sponge normally releases to control its pumping activity and causing an imbalance in the endogenous signaling feedback loops. Stimulation with glutamate causes dilation and then constriction of ECs, but at the end of the sneeze the excurrent canal diameter was usually smaller than when the sponges were at rest. This effect was more pronounced when sponges were treated with the higher concentration of 10 U/mL apyrase, and the higher enzymatic content also increased the variability of sponge responses (Fig. 3C). As with all physiological systems, sponges rely on a balance of signaling inputs and outputs to regulate their internal activities and maintain homeostasis. Our results suggest that ATP and glutamate work in the same pathway to coordinate contractions, but play different roles: ATP seems to cause the expansion of excurrent canals, whereas glutamate functions to constrict them.

### Contractions require ATP signaling

We then asked whether ATP is necessary for contractions or if it has a more modulatory role in contractions. Using 100 µM PPADS, a selective P2 receptor blocker (Lambrecht et al., 1992), we found that sponges did not show any response when stimulated with 70 µM glutamate or 100 µM ATP. This result strongly suggests that ATP is required to activate purinergic receptors in order to initiate a sneeze; it also confirms that ATP acts downstream of glutamate signaling.

### Proposed pathway for signaling by ATP and glutamate

Activation of P2X receptors triggers and modulates glutamate release (Gu and MacDermott, 1997). In neurons, antagonizing P2X receptors prevents enhanced glutamate signaling (Nakatsuka et al., 2003). Glutamate signaling in astrocytes causes the release of ATP which stimulates propagation of Ca^2+^ signals (Hamilton et al., 2008). We propose that a similar pathway regulates contraction behaviours in sponges as described in Fig. 5. A stimulus, either mechanical or chemical, is detected by sensory primary cilia lining the osculum, the chimney-like structure through which water is expelled from the sponge (Ludeman et al., 2014). These stimuli could result from clogging of the canals with sediments causing mechanical irritation or reduced flow through the sponge, or by intake of unwanted particles. The cilia in the osculum are analogous to the primary cilia of the kidney epithelium in function (Singla and Reiter, 2006): changes in water flow changes the bending of the cilia, triggering entry of Ca^2+^ into the cilium, and thereafter the release of signaling molecules such as glutamate from pinacocyte epithelia. Glutamate, acting through metabotropic glutamate receptors, then causes a cascade of downstream effects: the release of ATP, which acts in both an autocrine and paracrine manner to increase intracellular calcium via ionotropic P2X receptors and mobilization of internal calcium stores. This would then activate the contractile apparatus in cells, expanding the ECs by the contraction of ICs. Eventually, through the breakdown or reuptake of ATP, the cue for dilating the ECs diminishes while the glutamate-driven signal for constricting the ECs takes over.

**Figure 5.**
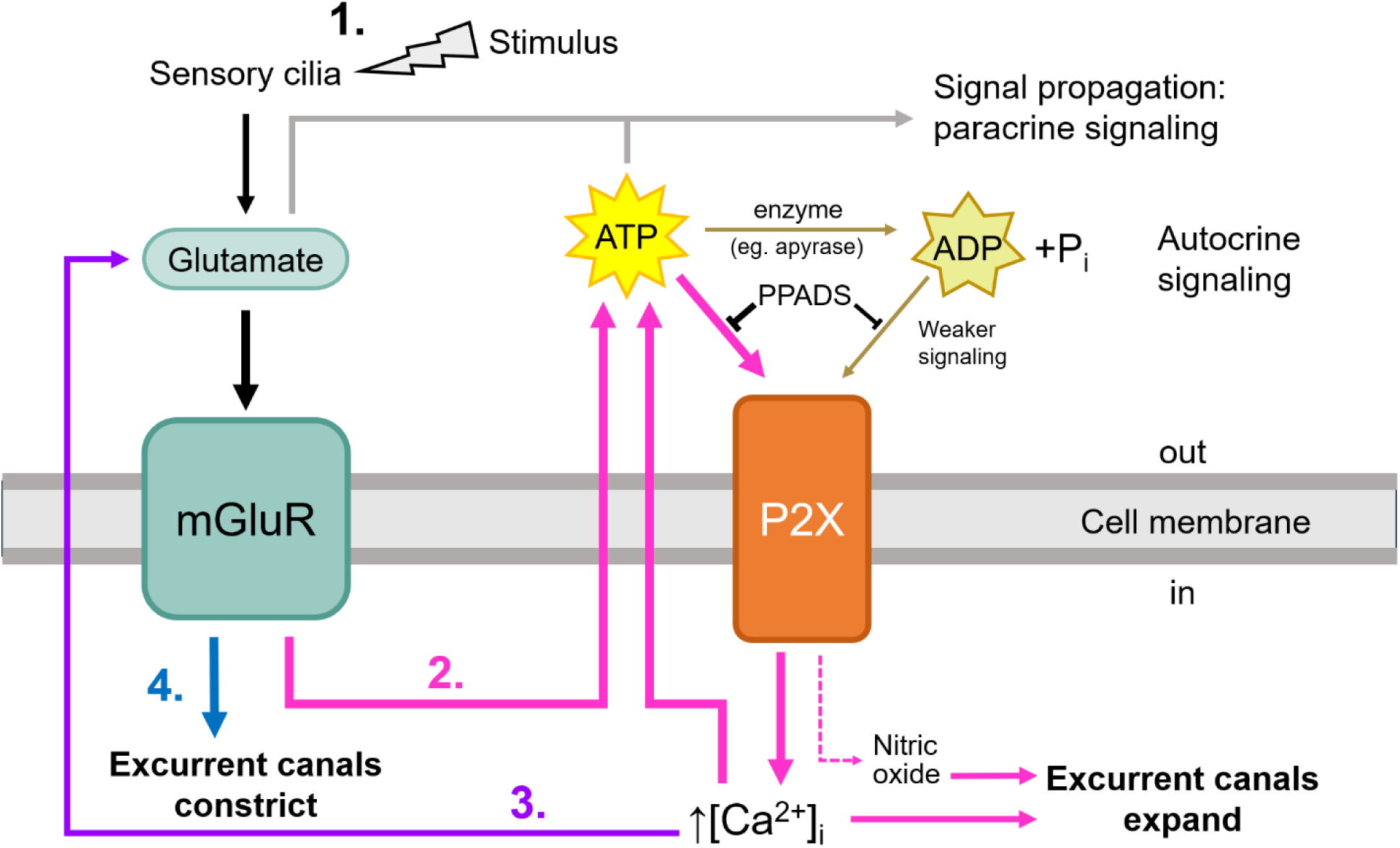
L-glutamate and ATP are key components of the signaling pathways involved in controlling contractions. A mechanical or chemical stimulus activates sensory cilia, triggering the release of glutamate. Metabotropic glutamate receptors (mGluRs) on contractile cells lining the aquiferous canal system become activated and trigger the release of ATP. ATP increases intracellular calcium levels and nitric oxide to expand the excurrent canals. The Ca^2+^ also aids in signal amplification and propagation by causing the release of more signaling molecules (e.g. ATP and glutamate). ATP and glutamate may work in both a paracrine and autocrine manner. Eventually, this signal for excurrent canal expansion will weaken as ATP is removed from the extracellular space and the signal for constricting the excurrent canals will take over, allowing the contraction to come to an end.

Despite blocking sneezes with PPADS, the pumping activity of the flagellated chambers did not stop. Therefore, it is likely that contractions are regulated separately from the sponge’s basal pumping activity, although this requires further investigation.

### P2X receptors are evolutionarily conserved

Sponge P2X receptor sequences share high sequence homology with those of well characterized P2X receptors. In fact, these receptors, though notably absent in insects and the nematode *Caenorhabditis elegans*, are highly conserved among most animal lineages (Fountain and Burnstock, 2009). P2X receptors themselves have been identified in non-metazoan eukaryotes including fungi, amoeba, and green algae (Fountain, 2013). It is plausible that P2X receptors had a key role in the advent of aggregating/colonial behaviours by mediating intercellular communication, and became integrated into a system of coordination in the first metazoans. As animals evolved increasing morphological complexity (e.g., defined organ systems), the role of P2X receptors also expanded. In vertebrates, several P2X receptor subtypes have been characterized, each with their own unique properties and roles in various physiological systems (North, 2016). Our results show that invertebrates also have several P2X receptor subtypes of their own. Some have high amino acid sequence similarity in the conserved motifs and functional domains as vertebrate P2X receptors, although invertebrate P2X receptors are not as well studied.

### Structural homology of P2X receptors

Phylogenetic analysis also revealed 2 distinct sets of receptors: one group that includes the vertebrate P2X receptors, and the other that appears to be exclusive to invertebrates. Whereas the set of vertebrate-included P2X receptors all contain the conserved amino acid sequence motifs that are characteristic of P2XRs, these sequence motifs are not present in the other set of receptor sequences (Supplemental data 3). The phylogenetic tree is rooted to the amoebozoan *D. discoideum*, the most evolutionarily distant organism from metazoa. Though the predicted phylogeny shows it has no relationship to other P2XRs, our multiple sequence alignment shows that 1 of 3 conserved motifs is present, along with other highly conserved domains. The presence of one of the conserved motifs in both a fungus (*G. margarita*) and an amoeba (*D. discoideum*) P2XR sequences but their absence from the invertebrate-only group suggests that the second group of P2XRs may be phylogenetically distinct purinergic receptors.

Further analysis of the receptors by 3D protein modeling shows that sponge P2XRs are nearly identical in structure to the zebrafish P2X4 subunit (Fig. 4C), of which the crystal structure is known and depicted as a ‘dolphin’ with extensive folding to create a ‘head’ region with the N- and C-terminal helices creating a ‘tail’ (Kawate et al., 2009). The structural similarity of the two *E. muelleri* P2X receptors suggests that while there is little amino acid sequence conservation, preserving the structure of the protein is more important to its function, which Leys and Riesgo (2011) also concluded when investigating type IV collagen in sponges.

Certain regions within the sequences are highly conserved among all our analyzed P2X receptors (Supplemental data 3). These are likely to be domains and specific residues that are critical for purinoceptor function. While P2X receptor subtypes themselves have different affinities for ATP (Hou and Cao, 2016), there is very little activation by ADP because of the positioning of positively charged residues within the ATP binding site (Chataigneau et al., 2013). Results from our physiological experiments showed that ADP was sufficient to trigger contractions in sponges (Fig. 3B). It is possible that receptors encoded by the Em0004g666a gene (and others in its group) are not as specific as conventional P2XRs for ligand binding. Differences in the sequence and structure may therefore allow activation by ADP, although this hypothesis requires further investigation.

## Conclusions

The present study demonstrates that sponges use purinergic signaling to coordinate behaviour. We provide evidence that ATP signaling is necessary for contractions in sponges to occur, and that sponges respond to ATP in a concentration-dependent manner. Interestingly, constant exposure to excess ATP in the bath application causes a sustained dilation of the excurrent canals, preventing the completion of the sneeze ‘expansion-contraction’ cycle, which suggest that normally there must be a mechanism for removing ATP from the extracellular space to prevent over-activation of the sponge tissues. Our finding that glutamate and ATP signaling work in concert to coordinate contractions of the sponge and the conservation of P2X receptors from sponges indicates that signaling pathways involved in coordinating sponge behaviour are highly similar to the signaling networks used in vertebrate central nervous systems.

## Table of abbreviations

ATP: Adenosine triphosphate
GABA: γ-amino-butyric acid
PPADS: pyridoxalphosphate-6-azophenyl-2’,4’-disulfonic acid
EC(s): Excurrent canal(s)
IC(s): Incurrent canal(s)
P2XR(s): P2X receptor(s)

## Acknowledgments

We gratefully acknowledge W. Gallin for assistance with the BLAST search of sponge transcriptomes and for fruitful insights on the molecular evolution of ion channels, and K. Guillas for many enjoyable conversations on the underlying physiology of sponge sneezes.

## Data availability

All relevant data can be found within the article and its supplementary information.

